# Plastic ingestion in Asian elephants in the forested landscapes of Uttarakhand, India

**DOI:** 10.1101/2020.12.14.422711

**Authors:** Gitanjali Katlam, Soumya Prasad, Anant Pande, Nirala Ramchiary

**Affiliations:** School of Life Sciences, Jawaharlal Nehru University, New Delhi – 110067, India; Nature Science Initiative, 36 Curzon Road, Dehradun- 248001, Uttarakhand, India; Wildlife Institute of India, Chandrabani, Dehradun- 248001, Uttarakhand, India

**Keywords:** Plastic pollution, elephant habitats, waste segregation, endangered species, terrestrial ecosystems

## Abstract

Impacts of plastic pollution, recognized as a driver of change in the global environment, have been under reported in terrestrial fauna. In this study, we looked at presence of plastic in the diet of Asian elephant and other megaherbivores in the forest habitats of Haridwar and Lansdowne, Uttarakhand state, India. We collected dung and pellet samples from forest edges and forest interiors and quantified plastic particles and other anthropogenic waste present. Each anthropogenic waste item was measured, weighed and sub-categorized into the type of plastic or other categories. Thirty-two percent of the elephant dung samples showed presence of plastic and other waste. Plastic particles comprised of 85% of the waste recovered from the dung with 100% occurrence in elephant dung samples (mean 47.08±12.85 particles per sample). We found twice as many plastic particles (85.27±33.7 per 100g of dung samples) in forest samples as compared to forest edge samples (35.34±11.14 plastic particles/100g of dung samples). Other non-biodegradable anthropogenic waste recovered from elephant dung (glass, metal, rubber bands, clay pottery and tile pieces) was found to be much higher for forest samples (34.79±28.41 items/100g sample) as compared to forest edge samples (9.44±1.91items/100g). This study is the first systematic documentation of occurrence of non-biodegradable waste in the diet of Asian elephants. Dominance of plastic compared to other non-biodegradable material in elephant dung samples highlights its widespread use and poor waste segregation practices. We recommend developing a comprehensive solid waste management strategy to mitigate the threat of plastic pollution around these critical elephant habitats.

## Introduction

Plastic pollution has been recognized as one of the major drivers of change in global environment, influencing ecosystem processes as well as human well-being (Hernandez-Gonzalez et al., 2018; Malizia & Monmany-Garzia, 2019). Plastic being difficult to degrade in the environment (Bin et al., 2020) has become ubiquitous in all ecosystems (Malizia & Monmany-Garzia, 2019; Townsend et al., 2019). Owing to extensive use of single-use plastic, poor disposal and lack of recycling, plastic particles have accumulated in terrestrial habitats (Barnes et al., 2009) including mountains, rivers, forests, oceans (Eriksen et al., 2014), within deep sea (Chiba et al., 2018), sea shores (Browne et al., 2011) and terrestrial habitats (Malizia & Monmany-Garzia, 2019).

Plastic pollution is known to impact > 650 marine species (UNEP, 2011) including zooplankton (Sun et al., 2017), crustaceans (Goldstein & Goodwin, 2013), fish (Lusher et al., 2013), sea turtles (Santos et al., 2015), seabirds (Trevail et al., 2015; Wilcox, et al. 2015) and marine mammals (Waluda & Staniland, 2013; Hernandez-Gonzalez et. al., 2018). Ecological impacts of plastic pollution are alarming as it causes physical injuries such as strangulation, movement restriction, amputations (Williams et al., 2011; Baulch and Perry, 2014; Sigler, 2014), internal injuries and starvation (Gall & Thompson, 2015), and even mortality (De Stephanis et al., 2013). Further, plastic pollution fosters biological invasions (Geyer et al., 2017; Malizia & Monmany-Garzia, 2019), transports chemical contaminants (Windsor et al., 2019) and poses grave threat to human health (Wilcox et al., 2015). Such pervasiveness of plastic pollution both in land and in ocean, may have long-lasting, distant and large-scale, cascading effect on ecological systems, defining its global change drivers’ characteristics (Malizia & Monmany-Garzia, 2019).

Impact of plastic pollution has been under-reported for terrestrial environments in comparison to marine environments (Malizia & Monmany-Garzia, 2019), especially in rivers, deep forests due to heterogenous distribution of plastics on land (Jambeck et al., 2015; Ng et al., 2018; Malizia & Monmany-Garzia, 2019). Though few recent studies have demonstrated its impacts on a variety of soil organisms (Liu et al., 2017; de Souza Machado et al., 2018a, 2018b) including earthworms (Lwanga et al., 2017) and snails (Panebianco et al., 2019), the effects on endangered terrestrial or freshwater fauna are comparatively less known (Holland, 2016; Blettler et al., 2018).

Given the lack of information on plastic pollution impacts on terrestrial fauna, we framed this study to ascertain the presence of plastic in the diet of Asian elephant (*Elephas maximus indicus*) in the forests of Uttarakhand state, India. In this region, Asian elephants inhabit human-modified habitats (Johnsingh et al., 1990; Williams et al., 2001) and thus come directly in contact with anthropogenic waste (Puri et.al., 2020). In this manuscript, we identified, characterized and quantified visible plastic and other anthropogenic waste in Asian elephants (faecal samples as proxy of ingestion) ranging in close proximity to human habitations. We determined if there is a difference between plastic presence in areas with high human presence compared to interiors of the forests and discuss its impacts on this wide-ranging, endangered species and its habitat.

## Methods

### Study area

This study was conducted in and around the forest habitats of Uttarakhand state of India. The intensive study sites included Laldhang, Gaindikhata and Shyampur villages near Haridwar forest division (30° 8’ to 29° 32’ N and 77° 42’ to 78° 22’ E) and Kotdwara town near Lansdowne forest division (30° 6’ to 29° 36’ N and 78° 18’ to 78° 43’ E). Gaindikhata (human population = 2817) and Shyampur (human population = 2472) are located close to a national highway (NH 34) while Laldhang (human population= 6896) lies at the edge of Haridwar forest division. Kotdwara is a highly populated town (human population = 1,75,232) situated adjoining Lansdowne forest division (Figure 1).

**Figure 1.**
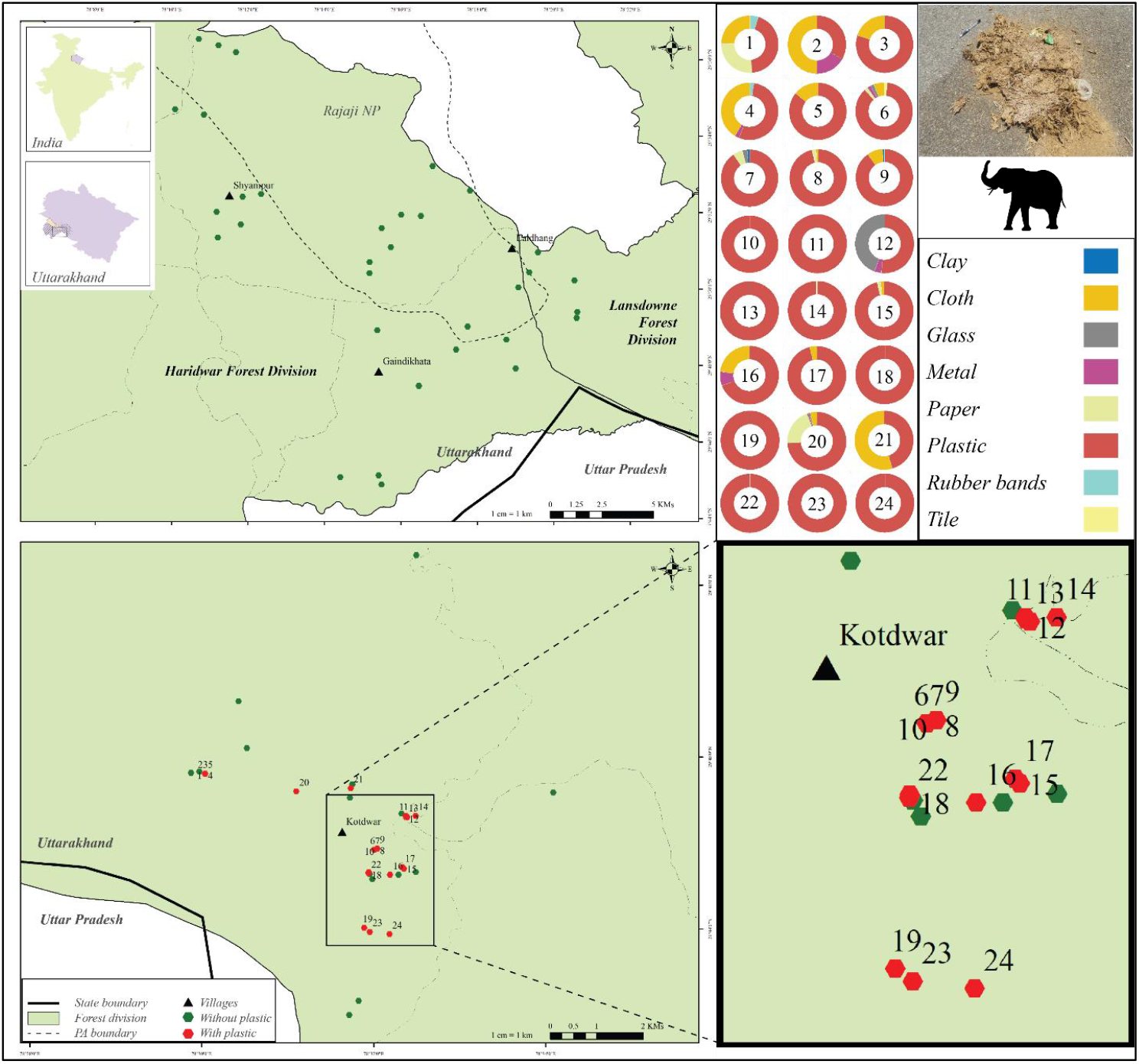
Study sites of Haridwar Forest Division (top left) and Lansdowne Forest Division (bottom left) adjacent to Rajaji National Park. Red and green dots represent sampling locations of elephant dung samples with and without plastic, respectively. Doughnuts (top right) represent proportion of plastic and other anthropogenic waste items retrieved from Asian elephant dung samples collected in and around Lansdowne Forest Division (bottom right).

These study sites consist of a mosaic of agropastoral land interspersed with dry seasonal river streams, open and mixed plantations, human habitation and road networks adjoining protected forest habitats. They are characterized by tropical dry and moist deciduous forests, dense shrub undergrowth and grassland habitats with high annual precipitation (1000 - 2500 mm / annum; Chitale, 2014). Vegetation at these sites is categorized as miscellaneous forest comprising of *Shorea robusta* mixed with *Mallotus philippensis, Ehretia laevis, Lagerstroemia parviflora, Albizia lebbeck, Azadirachta indica, Butea monosperma, Bauhinia purpurea, Adina cardifolia* etc. Dense shrub vegetation is dominated by invasive growth of *Lantana camara, Cassia tora, Parthenium hysterophorus* mixed with native species of *Justicia adhatoda, Murraya koenigii, Colebrookea oppositifolia, Ziziphus mauritiana* etc. Pure stands of *Shorea robusta* dominate inside protected forest areas whereas monoculture plantations (*Tectona grandis, Dalbergia sissoo* and *Eucalyptus* sp.) with mixed vegetation exists outside forest areas.

These study sites are part of north-western Terai-Arc Landscape, an important landscape for conservation of several threatened species such as tiger *Panthera tigris*, leopard *Panthera pardus*, northern swamp deer *Rucervus duvaucelii duvaucelii* and Asian elephant *Elephas maximus* (Johnsingh & Negi, 2003; Joshi, 2016; Paul et al., 2020). This landscape holds three Protected Areas i.e., Rajaji National Park, Corbett National Park and Jhilmil Jheel Conservation Reserve, amidst a mosaic of non-protected forest habitats and dense human habitations signifying its conservation importance. Haridwar and Lansdowne forest divisions act as immediate buffers of the Rajaji-Corbett Tiger Conservation Unit (Johnsingh & Negi, 2003) and constitute elephant corridors of high ecological priority (Tiwari et al., 2017). However, rapidly expanding human population, increasing road traffic and fragmentation of these migratory corridors over last couple of decades has aggravated the threat to native wildlife species especially along the Laldhang-Kotdwara forest habitats.

### Field sampling

Transects were sampled inside forest areas for fecal samples of elephants and other wild herbivores. In forest edges and villages, we searched and sampled opportunistically to locate elephant fecal samples (as it’s rare to find them outside the forest area). All sampling was carried out in the dry season between February to June 2018. The transects, 1 to 3 Km in length and spaced from each other by at least 2 Km, were laid starting from or nearby garbage dumps (at forest edges) towards the interior of the forest area. All the transects were sampled once during the field season.

### Collection of fecal samples

Dung samples of Asian elephant and pellet samples of other herbivores viz. barking deer (*Muntiacus muntjak*), nilgai (*Boselaphus tragocamelus*) and sambar *Rusa unicolor* were collected. Samples were hand-picked using sterile nitrile gloves in a beaker of 250 ml volume and kept in sterilized zip lock bags. Up to 4 sub-samples (each 250 ml volume) were collected from each elephant dung bolus encountered during surveys. All the dung/pellet samples were air dried and stored in sterilized zip lock bags labelled with date of collection, species, geographic location and site information (block name, forest range). The samples were later brought to the laboratory at School of Life Sciences, Jawaharlal Nehru University, New Delhi for further processing.

### Sample processing

Standardized protocols were used for sorting and quantification of anthropogenic wastes in a contamination-free laboratory environment (Van Franker et al., 2002; Klare et al., 2011; Hernandez-Gonzalez et.al., 2018). The work area and tools were sanitized before and after use. Samples were handled with sterilized nitrile gloves wearing cotton lab coats. All equipment (forceps, petri dishes, beakers) were cleaned thoroughly between samples using filtered water and absolute ethanol. Beakers containing samples were kept covered with aluminium foil to avoid any contamination. Each sub-sample was weighed on a fresh aluminum foil with the aid of an electronic balance (Citizon, max = 300g, d=10 mg). Tightly compacted dung boluses/pellets were carefully loosened up with forceps, measured and anthropogenic wastes visible to the naked eye were separated from the sample.

Total number of plastic particles and other anthropogenic waste was counted from each sub-sample visible to eye (>1mm), measured for length where widest, or diameter for circular ones and weighed them on an electronic balance (accuracy = 0.01 g). Visible plastic particles were further sub-categorized as disposable cutlery pieces, plastic pieces, plastic packaging and polythene bags. Further, plastic items were size classified as macroplastic (> 5 mm) and microplastic (1 - 5 mm) visible to naked eye (Di-Meglio et al., 2017; Hernandez-Gonzalez et al. 2018). The other anthropogenic waste was also categorized as non-biodegradable and biodegradable waste.

### Data analysis

All opportunistic dung/pellet samples collected along the forest edge and dung/pellet samples found within 100 m from the forest edge on transects were considered as forest edge samples. Similarly, dung/pellet samples collected from more than 100 meter from the forest edge up to 3 Km inside the forest during transect surveys, were considered as forest samples. Plastic and other waste for which length, weight and width could not be measured were not considered for the analysis. Overall, data is presented as mean abundance with standard error. All analyses were performed using the R program using packages “ggplot2” (Wickham, 2016), “beanplot” (Kampstra, 2008) “plotly” (Sievert, 2020) in R program v. 3.6.0 (R core team, 2019).

## Results

We conducted a total of 26 transects with survey effort of 68.2 Kilometer across the four blocks covering a total area of ~ 273 Km^2^ (Figure 1). Plastic particles and other anthropogenic waste were retrieved from 32% of the elephant fecal samples all belonging to sampling sites in Kotdwara area (24 samples - 14 forest edge and 10 forest). Overall, 75 elephant dung samples were collected during transects (n=64) and opportunistic (n=11) sampling from Kotdwara (40), Laldhang (11), Shyampur (18) and Gaindikhata (6). We did not find any plastic or any anthropogenic waste visible to naked eye in the fecal samples of sambar (n = 69), barking deer (n = 7) and nilgai (n = 56).

### Composition and abundance of plastic particles in elephant dung

We retrieved a total of 1130 plastic particles from 24 elephant dung samples (Figure 2; Supplementary Figure 1). Plastic particles comprised of 85% of the waste recovered from the dung with 100% occurrence in elephant fecal samples; ranging from 1 to 220 plastic particles per sample (Table 1). Disposable cutlery pieces (47.75±8.7 particles/sample) and plastic pieces (25.15±8.51 particles/sample) made up the most frequent plastic items, followed by plastic packaging (4.18±1.25 particles/sample) and polythene bags (1.6±0.18 particles/sample) (Figure 2). Overall mean abundance of plastic particles in elephant dung samples was estimated to be 47.08±12.85 particles per sample. In forest samples, higher abundance of plastic particles per sample were recorded (74.3±22.88 particles/sample) in comparison to forest edge samples (27.64±8.29 particles per sample, (Wilcoxon test, W=54.5, p> 0.05; Figure 3 a). Macroplastics (38.33±10.09 particles/sample) were observed to be more abundant as compared to microplastics (11.85±3.23 particles/sample) (Figure 4). In forest samples, count for plastic particle was recorded as 85.27±33.7 per 100g of dung samples, which is more than twice as compared to forest edge samples (35.34±11.14 plastic particles/100g; Figure 3c), in terms of weight 11.21±3.26 g of plastic particles/100g in forest samples were observed as compared to forest edge samples (3.7±0.72g /100g, χ^2^ = 20.062, df=1, p< 7.497e-06; Figure 5.3e). Higher incidence of plastic particle in dung were recorded from sampling sites in Totgadhera beat (in abundance - 166.57±199 particles/sample) and Lalpani beat (in weight - 0.54 ±0.8 g/ 100g of elephant dung; see Table 2).

**Figure 2.**
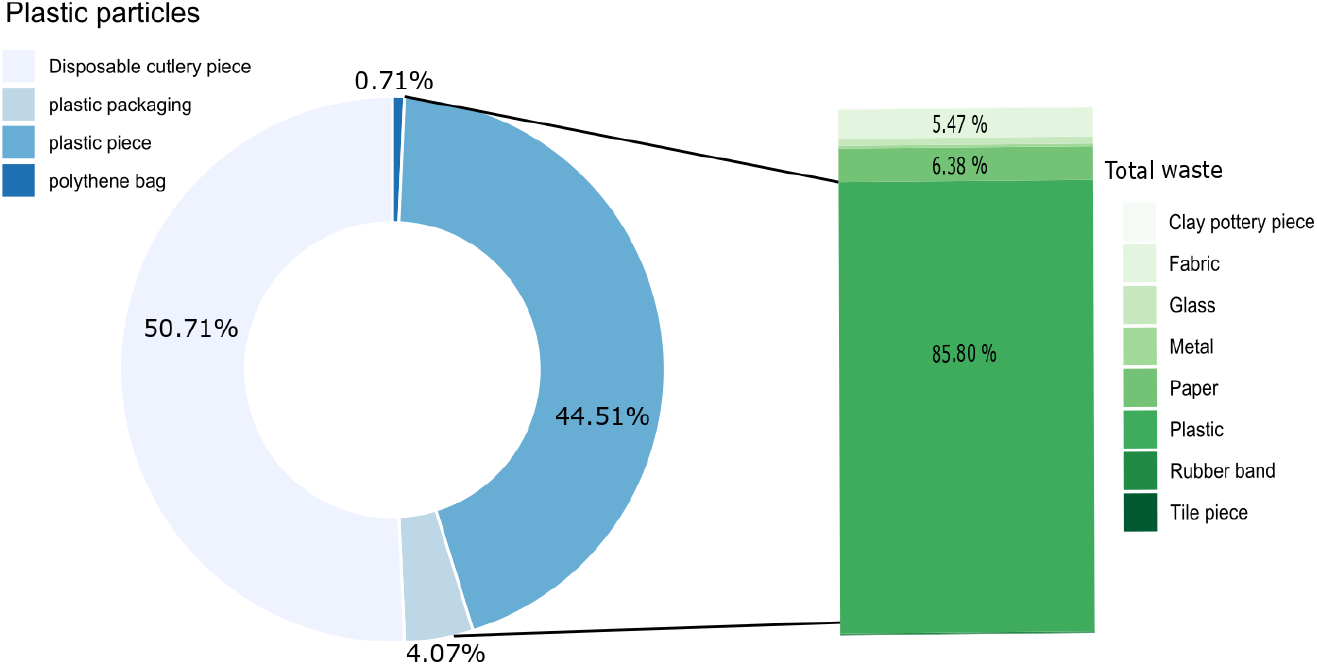
Percentage composition of plastic particles and total anthropogenic waste items retrieved from Asian elephant *Elephas maximus* dung samples collected from in and around Lansdowne Forest Division, Uttarakhand, India.

**Table 1.**
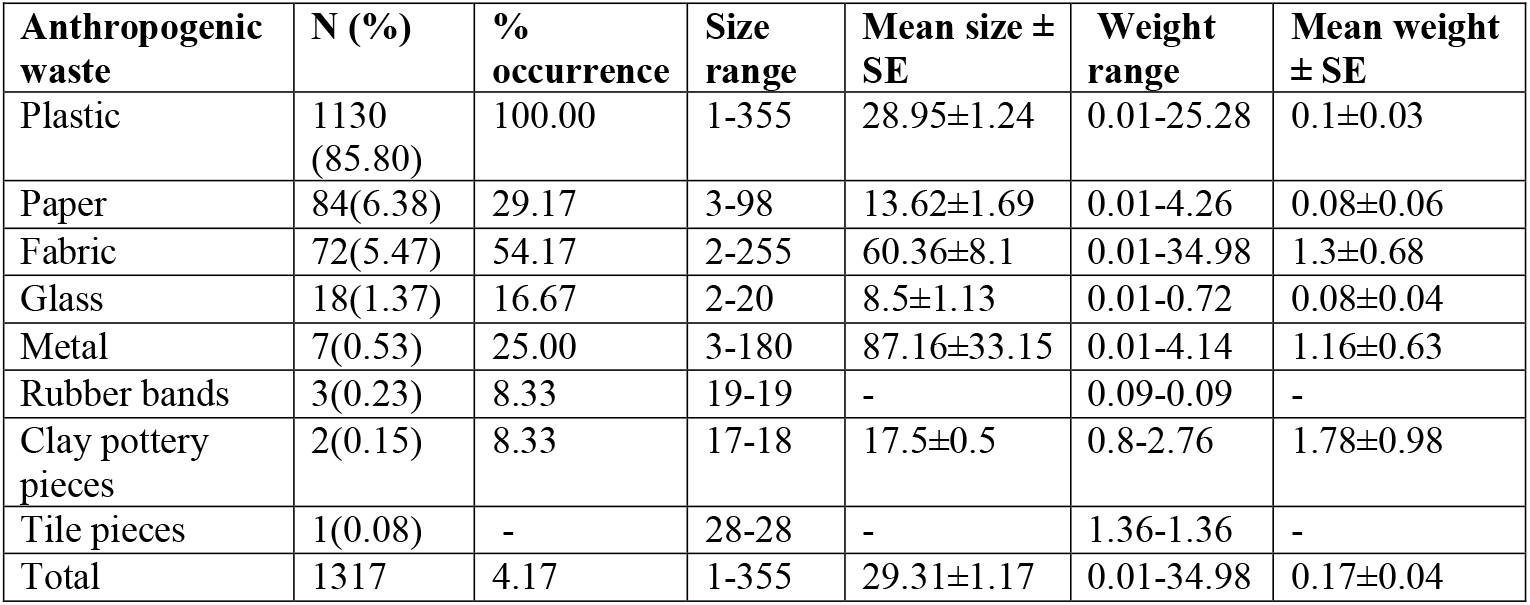
Anthropogenic waste recovered and identified from Asian elephant dung samples collected in and around Lansdowne forest division, Uttarakhand.

**Figure 3.**
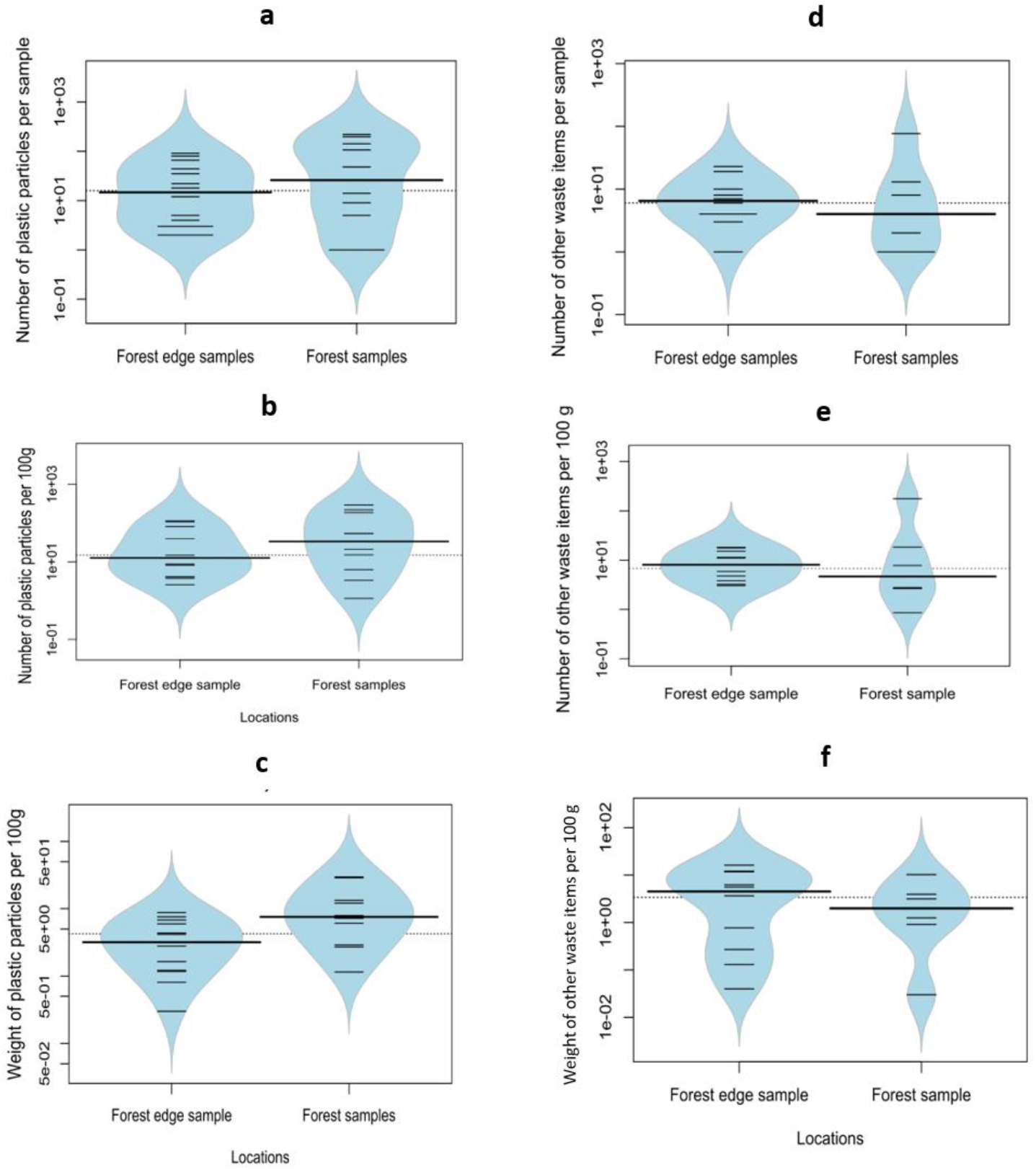
Bean plots depicting a) mean abundance of plastic particles per sample; b) mean abundance of plastic particles per 100 grams of sample; (c) mean weight of plastic particles per 100 grams of sample; (d) mean abundance of anthropogenic wastes per sample; (f) mean abundance of other waste items per 100 grams of sample; and (f) mean weight of other waste items per 100 grams of sample retrieved from Asian elephant dung samples collected in and around Lansdowne Forest Division, Uttarakhand, India.

**Figure 4.**
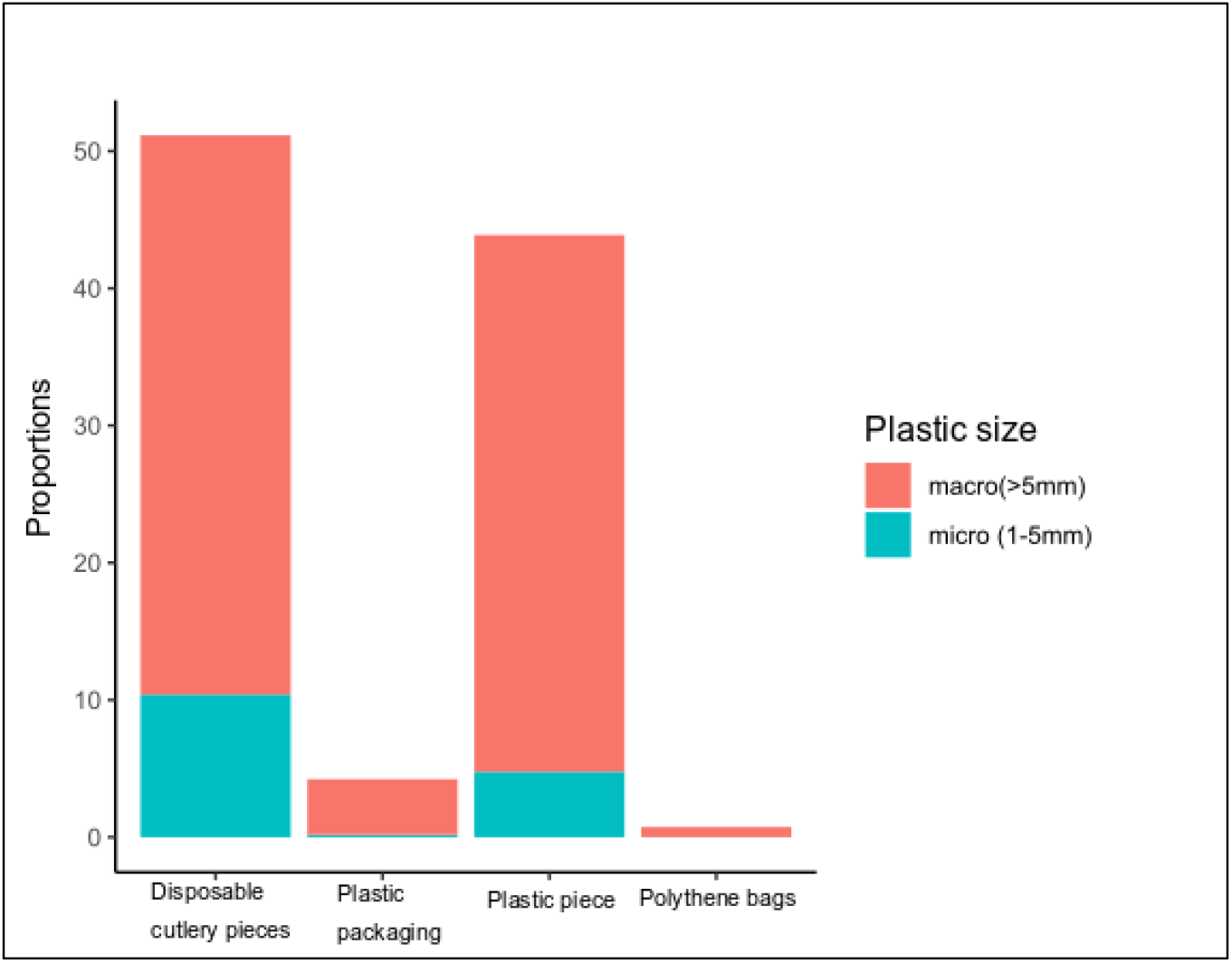
Percentage composition of macroplastic and microplastic particles in Asian elephant dung samples collected in and around Lansdowne Forest Division, Uttarakhand, India.

**Table 2.**
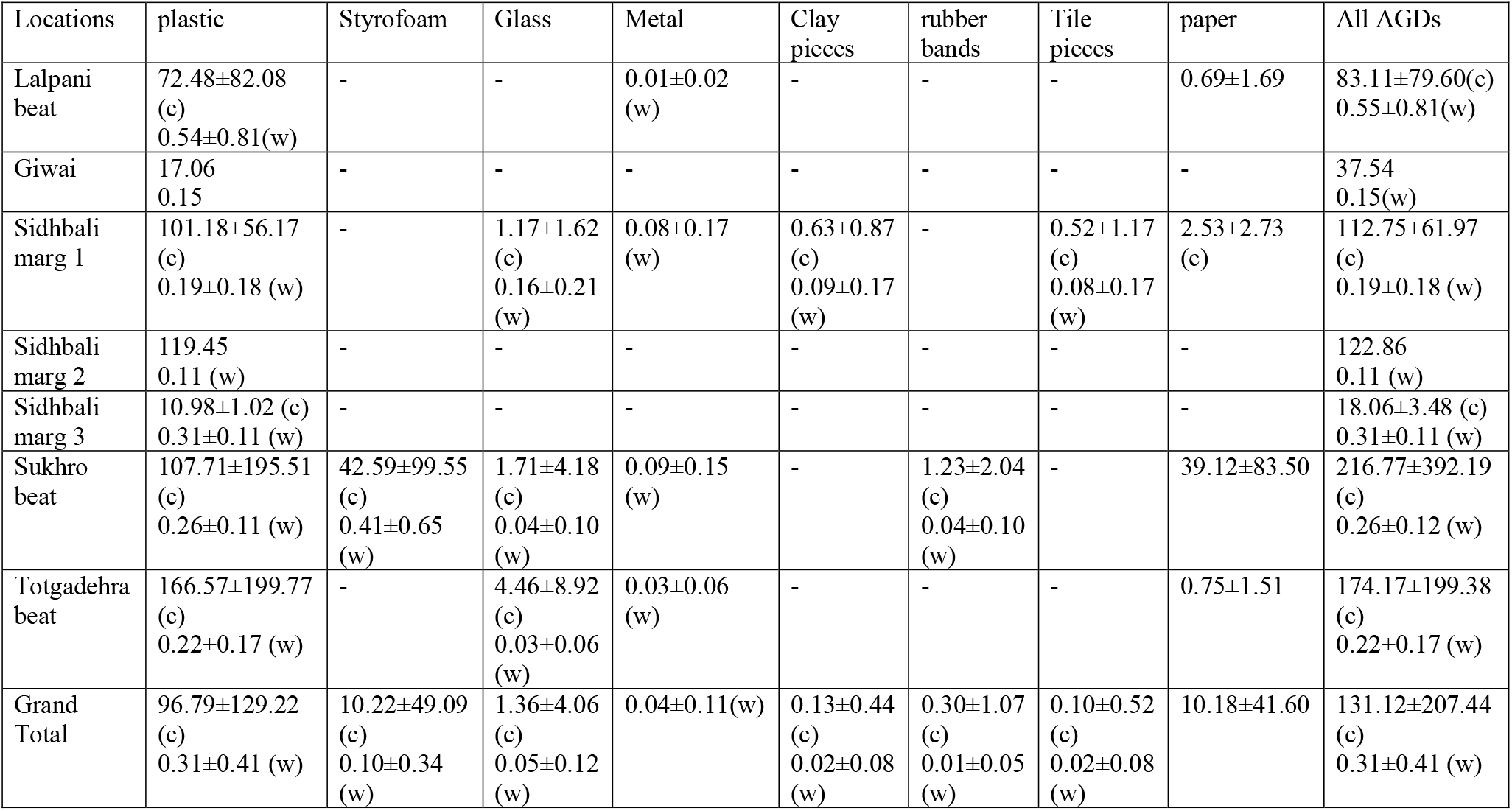
Mean count (c) and weight (w) of anthropogenic debris (AGD) items per 100g of elephant dung samples of Kotdwara study site, Uttarakhand, India.

| Locations | plastic | Styrofoam | Glass | Metal | Clay pieces | rubber bands | Tile pieces | paper | All AGDs |
| --- | --- | --- | --- | --- | --- | --- | --- | --- | --- |
| Lalpani beat | 72.48±82.08 (c)<br>0.54±0.81(w) | - | - | 0.01±0.02 (w) | - | - | - | 0.69±1.69 | 83.11±79.60(c)<br>0.55±0.81(w) |
| Giwai | 17.06<br>0.15 | - | - | - | - | - | - | - | 37.54<br>0.15(w) |
| Sidhbali marg 1 | 101.18±56.17 (c)<br>0.19±0.18 (w) | - | 1.17±1.62 (c)<br>0.16±0.21 (w) | 0.08±0.17 (w) | 0.63±0.87 (c)<br>0.09±0.17 (w) | - | 0.52±1.17 (c)<br>0.08±0.17 (w) | 2.53±2.73 (c) | 112.75±61.97 (c)<br>0.19±0.18 (w) |
| Sidhbali marg 2 | 119.45<br>0.11 (w) | - | - | - | - | - | - | - | 122.86<br>0.11 (w) |
| Sidhbali marg 3 | 10.98±1.02 (c)<br>0.31±0.11 (w) | - | - | - | - | - | - | - | 18.06±3.48 (c)<br>0.31±0.11 (w) |
| Sukhro beat | 107.71±195.51 (c)<br>0.26±0.11 (w) | 42.59±99.55 (c)<br>0.41±0.65 (w) | 1.71±4.18 (c)<br>0.04±0.10 (w) | 0.09±0.15 (w) | - | 1.23±2.04 (c)<br>0.04±0.10 (w) | - | 39.12±83.50 | 216.77±392.19 (c)<br>0.26±0.12 (w) |
| Totgadehra beat | 166.57±199.77 (c)<br>0.22±0.17 (w) | - | 4.46±8.92 (c)<br>0.03±0.06 (w) | 0.03±0.06 (w) | - | - | - | 0.75±1.51 | 174.17±199.38 (c)<br>0.22±0.17 (w) |
| Grand Total | 96.79±129.22 (c)<br>0.31±0.41 (w) | 10.22±49.09 (c)<br>0.10±0.34 (w) | 1.36±4.06 (c)<br>0.05±0.12 (w) | 0.04±0.11(w) | 0.13±0.44 (c)<br>0.02±0.08 (w) | 0.30±1.07 (c)<br>0.01±0.05 (w) | 0.10±0.52 (c)<br>0.02±0.08 (w) | 10.18±41.60 | 131.12±207.44 (c)<br>0.31±0.41 (w) |

### Composition and abundance of other anthropogenic waste in elephant dung

Other anthropogenic waste recovered from elephant dung consisted of non-biodegradable wastes such as glass (n= 18) pieces > metal (n = 7) > rubber bands (n = 3)> clay pottery (n = 2) and tile (n = 1) pieces (Supplementary Figure 2). The biodegradable anthropogenic waste recovered from samples consisted of paper (n=84), fabric pieces (n=72), and human hair fragments (n=5) (Figure 2, Table 1).

The overall mean abundance of other waste was observed to be 11.24±4.38 items per sample. The forest samples again showed higher abundance of these (26.5±9.96 items/sample) as compared to forest edge samples (15.45±5.83 items/sample) (Wilcoxon test, W=40.5, p> 0.05; Figure 3b). The mean count for other waste per 100g of total dung sample was found to be higher in forest samples (34.79±28.41 items/100g sample) as compared to forest edge samples (9.44±1.91items/100g; Figure 3d). Similarly, the mean weight of other waste per 100g was found to be more or less similar in forest samples (3.24±1.51g/100g) and forest edged samples (5.66±1.85g/100g, χ^2^ =16, df=15, p>0.05; Figure 3f).

## Discussion

To our knowledge, this study is the first systematic documentation of occurrence of non-biodegradable waste, plastic particles, other hazardous and toxic anthropogenic waste in the diet of Asian elephants. We retrieved plastic particles from elephant dung samples which were collected from Kotdwara town, where a large human population lives in close proximity of the forest (Census of India, 2011). Dominance of plastic compared to other non-biodegradable material in the Kotdwara elephant dung samples indicates its widespread use (due to low-cost availability - Derraik, 2002) and poor waste management in the area. The occurrence of other hazardous material (metal, glass, cloth fabric) in the dung samples highlights poor waste segregation practices despite a higher-than-average literacy rate (~80%) in the area (Census of India, 2011).

Asian elephants were found to forage near forest edges on garbage dumps carrying food waste (Puri et. al, 2020) and ingest plastic mixed with other non-biodegradable waste. We found high occurrence of macroplastic particles in the elephant dung (mostly disposable cutlery and polythene bags, plastic packaging), seemingly influenced by foraging behaviour of elephants. As gulpers (Katlam et.al., 2018), elephants are likely to ingest large portions of food waste mixed with plastics and other hazardous waste material. We found more than twice the number of plastic particles in forest samples as compared to forest edge samples, signifying ingression of plastic particles into forest areas through elephants. These deposited plastics might degrade into smaller particles and transfer through trophic invertebrates’ prey to predators such as birds (D’Souza et al., 2020) with potentially negative impacts. Further, the plastic particles may spread far and wide into the forest systems away from human presence as elephants can move several kilometers in a day depositing dungs (Williams et al., 2001).

Rajaji-Corbett landscape suffers from habitat fragmentation leading to mosaic landscapes with poor to loss of connectivity between forest patches (William et al., 2001; Johnsingh et al., 2004). Increased diversion of forest land, overgrazing, excessive lopping of trees for forage, and infrastructure development in the region thus threatens the extant elephant population of the region (William et al., 2001). Our study highlights emergence of a new threat in the form of plastic pollution to endangered Asian elephants with increasing human occupation of the forest edges around this landscape. Plastic ingestion by elephants and other species visiting garbage dumps would not only be detrimental to individuals but to forest ecosystems with an impact on lower trophic levels (D’Souza et al., 2020; Jâms et al., 2020) through animal-aided dispersal. Overall, our data demonstrates the negative impacts of improper waste management on an endangered species around protected areas of conservation significance. We recommend developing a comprehensive solid waste management strategy through mapping of garbage dumps, conducting risk assessment to the wildlife and mass awareness campaigns to mitigate the threat of plastic pollution around these critical elephant habitats.

## Acknowledgments

The financial support for this work was provided to Gitanjali Katlam by The Rufford Foundation, United Kingdom (Grant ID: 19961-1). Logistics at field was supported by Nature Science Initiative, Dehradun. Authors are thankful to Dean, School of Life Sciences, Jawaharlal Nehru University, New Delhi and Chief Wildlife Warden, State Forest Department, Uttarakhand for providing necessary permissions to carry out laboratory analysis and field work, respectively. We are grateful to the field assistants, Basheer, Zareef, Shamshad, Mumtaz, Saddam, Taukir, Neel, Shreyash, Sumit and Lokesh for help during field work and Sohom for help in preparation of the maps.

---

**Supplementary Figure 1.**
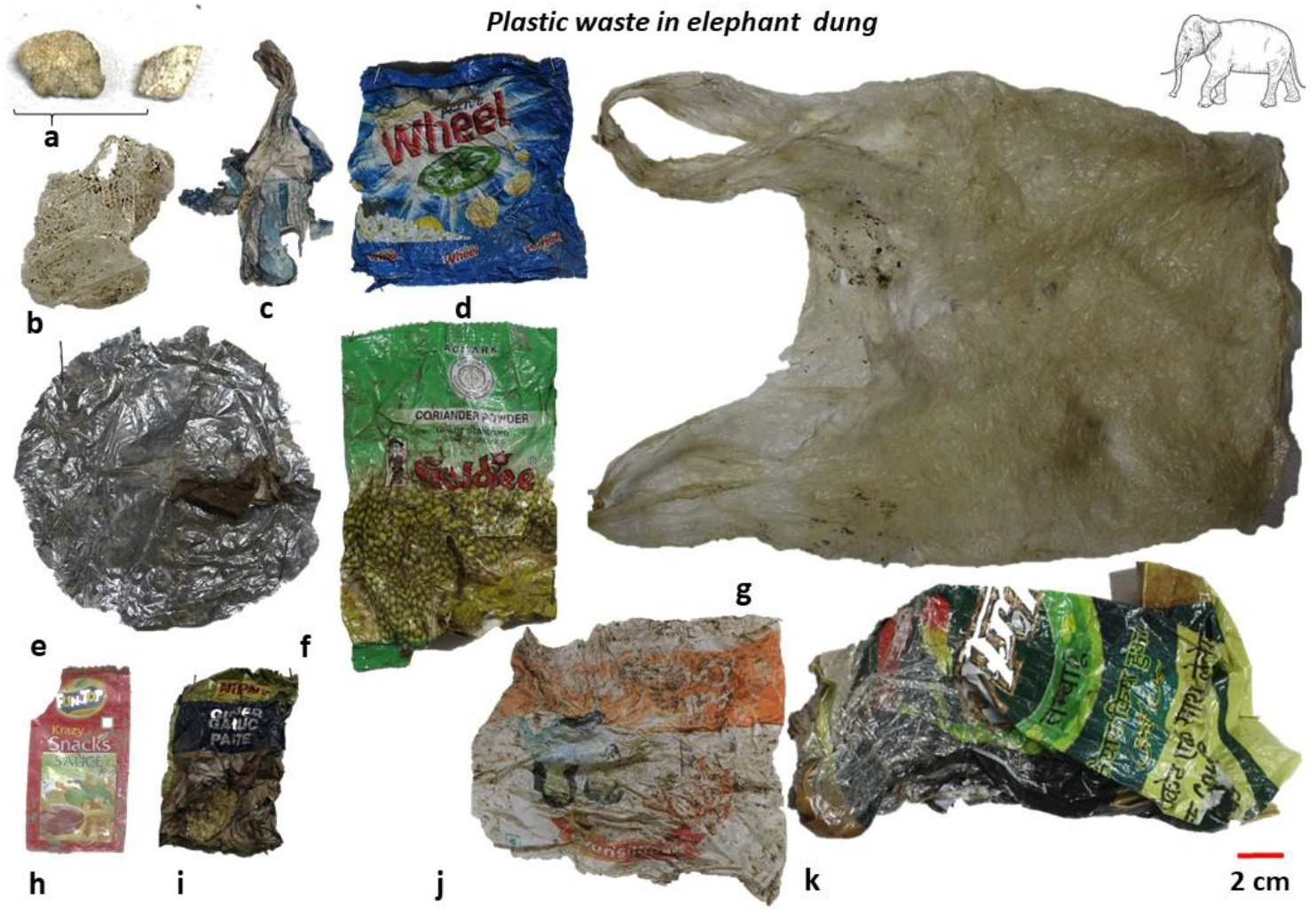
Types of plastic items retrieved from Asian elephant dung samples collected in and around Lansdowne Forest Division, Uttarakhand, India. a) styrofoam, b) disposable plastic cup, c) plastic tube, d) detergent packaging, e) disposable plate, f) spice powder packaging, g) polythene bag, h) ketchup sachet, i) spice paste packaging, j) milk packet and k) tobacco packaging.

**Supplementary Figure 2.**
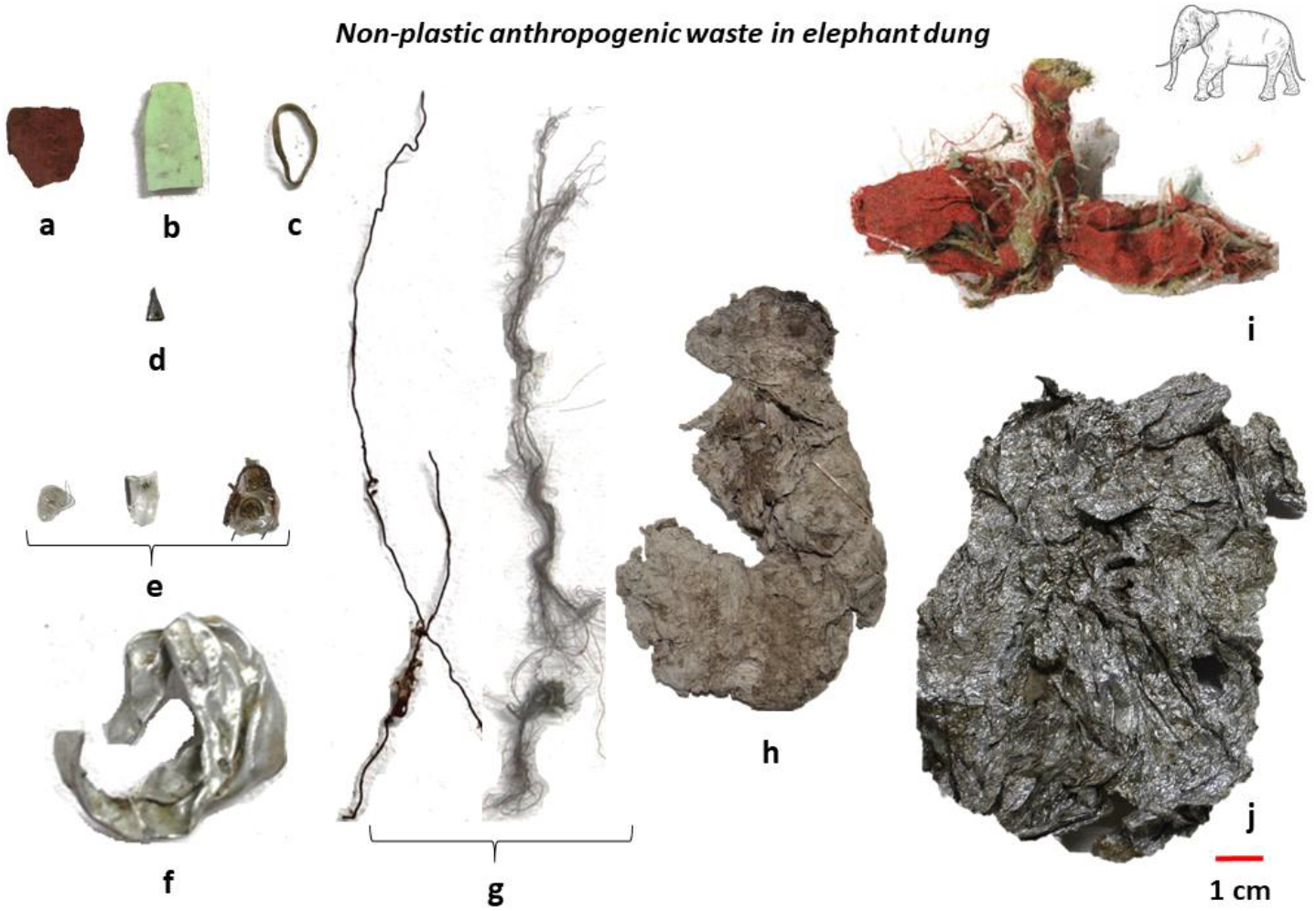
Types of non-plastic waste items retrieved from Asian elephant dung samples collected in and around Lansdowne Forest Division, Uttarakhand, India. a) clay pottery, b) tile piece, c) rubber band, d) glass piece, e) pieces of filament bulb, f) metal screw base of a filament bulb, g) metal wires, h) ketchup sachet, i) synthetic fabric, and j) aluminium foil.

